# SARS-CoV-2 and other coronaviruses in rats, Berlin, Germany, 2023

**DOI:** 10.1101/2023.12.15.571826

**Authors:** Kerstin Wernike, Calvin Mehl, Andrea Aebischer, Mario Heising, Rainer G. Ulrich, Martin Beer

## Abstract

We tested 130 rats trapped in Berlin for coronaviruses. Antibodies against SARS-CoV-2 were detected in a single animal only, but not in further 66 rats from the same location, speaking against virus circulation in the rat population. All animals tested negative for SARS-CoV-2 by RT-PCR. However, rodent-associated alphacoronaviruses were found.

## Main text

The severe acute respiratory syndrome coronavirus 2 (SARS-CoV-2), a betacoronavirus, was initially reported in 2019 in China and thereafter spread rapidly worldwide, causing the COVID-19 pandemic in humans. Since the pandemic unfolded, it was speculated about the role of animals as amplifying or reservoir hosts. Because of the long-term association between rodents and coronaviruses *(1)*, the wide range of coronaviruses occurring in wild rodents *(2)* and the ubiquitous distribution of commensal rodents, it was obvious to also include rodents in susceptibility studies, among them rats. Under experimental conditions using high infection doses, rats were reported as receptive particularly to the SARS-CoV-2 delta variant of concern (VOC), but also experimental infection with other variants like alpha, beta or omicron were described *(3,4)*, posing the theoretical risk for establishing effective infection chains in nature. Accordingly, field studies were initiated early into the pandemic to investigate the situation in wild rats. Indeed, serological and molecular evidences of SARS-CoV-2 infection of a few animals could be found in some studies *(2,3,5)*, while others reported consistently negative results *(6,7)*. However, these studies were conducted before the emergence and worldwide large-scale spread of the omicron VOC and its diverse subvariants. In laboratory settings, lungs from omicron-infected animals showed significantly lower infectious viral titers compared e.g. to delta *(3)*, but field studies about omicron occurrence in rat populations are missing. Therefore, we investigated rats trapped in Berlin, the very densely populated (>4,000 inhabitants per km^2^) capital of Germany, during 2023, i.e. a period at which omicron represented the dominant variant in the human population.

Lung and chest cavity lavage fluid samples were collected from 130 Norway or brown rats (*Rattus norvegicus*) caught in the context of pest control at 44 trapping sites within Berlin (Figure 1A). The lavage fluids were tested for antibodies against SARS-CoV-2 by a receptor binding domain (RBD)-based multispecies ELISA using a cut-off of ≥0.3 for positivity as described *(8)*. Two orthologs of the RBD protein were used in parallel, the wild-type virus RBD and that of the omicron XBB1.5 variant. The samples were prediluted 1/10 as described for lavage samples of rodents *(6)*. One of the 130 rats tested positive, the optical density (OD) values were 1.16 (wild-type RBD) and 1.53 (omicron RBD), respectively. To confirm the positive results, the sample was additionally tested by a surrogate virus neutralization test (sVNT) (cPass SARS-CoV-2 Neutralization Antibody Detection Kit, GenScript, the Netherlands) performed as prescribed by the manufacturer (cut-off for positivity at ≥30% inhibition). In its original composition, the test enables the detection of antibodies against the wild-type virus and all VOCs except omicron. For omicron and its sub-variants, a specific RBD is provided by the test manufacturer. The rat sample positive in the RBD-ELISA was analyzed by the sVNT using the original and the omicron-specific RBD, and the omicron-based test gave a positive result (33.9% inhibition; 23.4% for the wild-type RBD). These results hint at a previous infection of the animal with an omicron subvariant. However, that only one rat tested positive speaks very clearly for a single spillover event from the human into the rat population and against autonomous virus circulation in rats, especially as further 66 rats were caught in the same building as the sero-reactive animal and all of them tested negative (Figure 1A).

**Figure 1.**
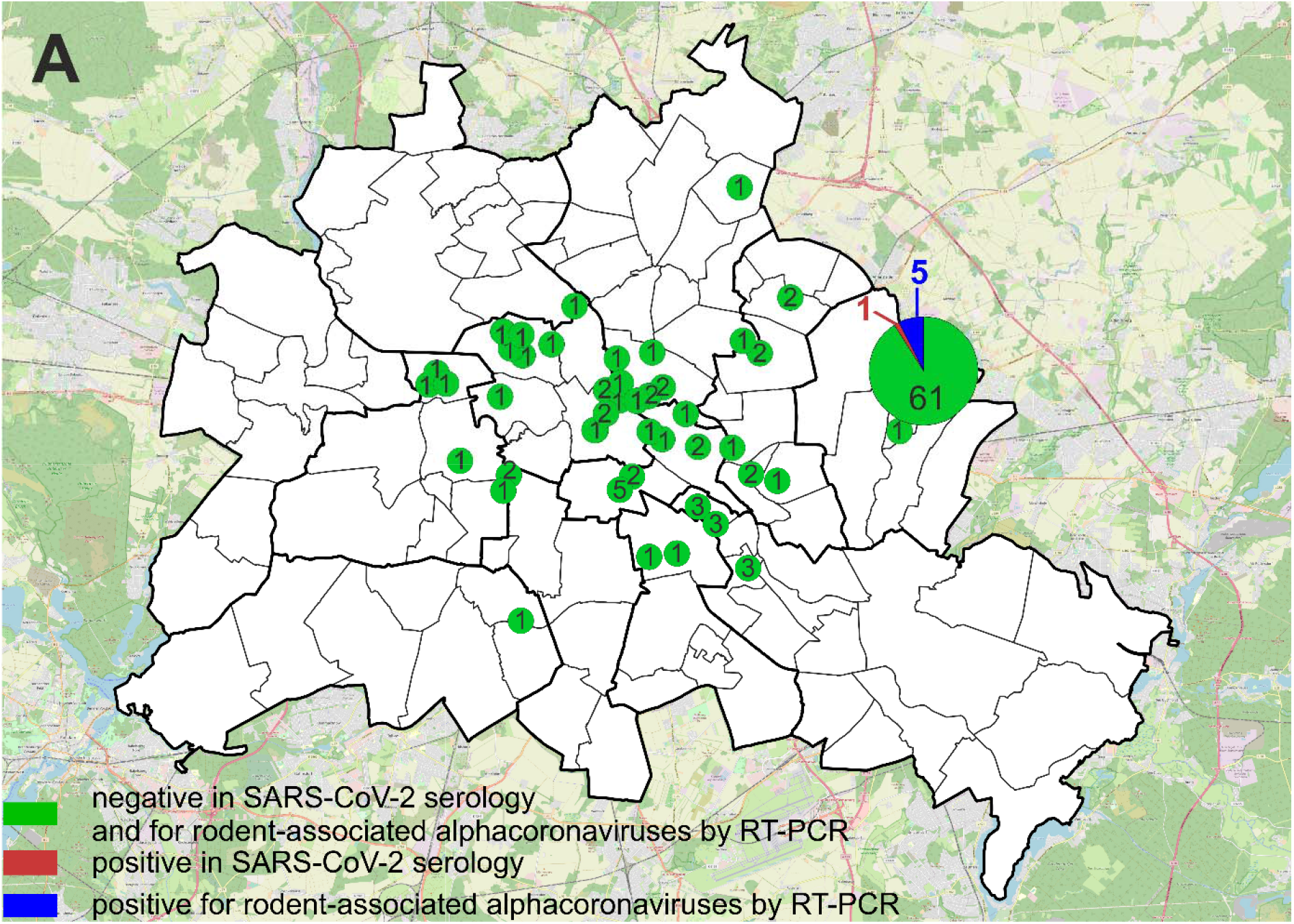
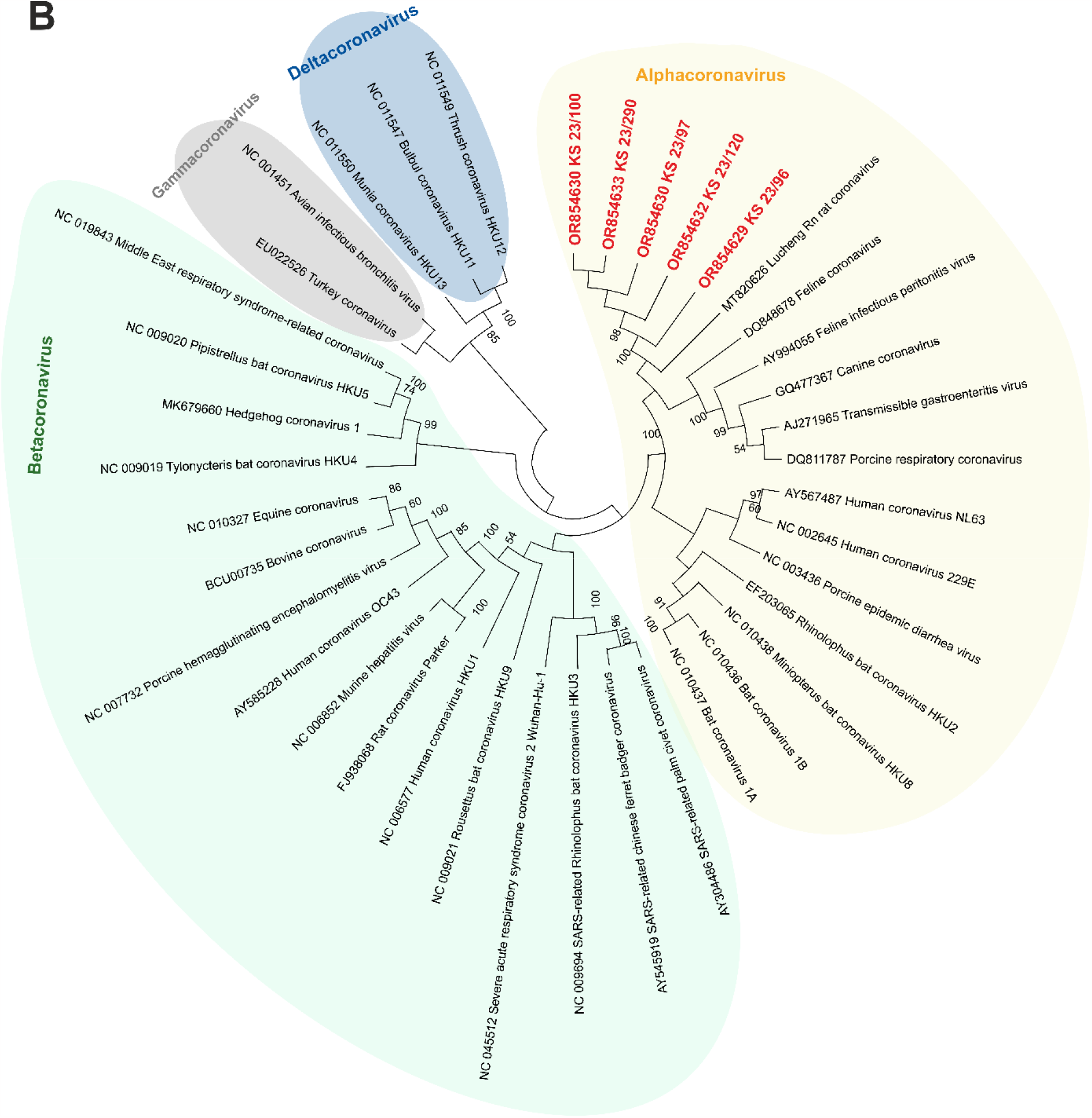
**A**. Locations within Berlin at which rats were trapped and number of animals per location. Green dots represent areas from which all sampled animals tested negative in the SARS-CoV-2 RBD-based ELISA and negative for coronaviruses by RT-PCR. When rats tested positive for coronaviruses by RT-PCR, the number of animals is given in blue. The single animal that tested positive for antibodies against SARS-CoV-2 is indicated in red. The map of Berlin, in which the dots were printed, was retrieved from Geoportal Berlin, dataset “Geoportal Berlin / Ortsteile von Berlin”, URL: https://daten.odis-berlin.de/de/dataset/ortsteile/, data license Germany – attribution – Version 2.0 (www.govdata.de/dl-de/by-2-0). The map of the area surrounding Berlin was retrieved from OpenStreetMap (map data copyrighted OpenStreetMap contributors and available from https://www.openstreetmap.org). **B**. Classification of the detected coronaviruses based on partial sequences of the RNA-dependent RNA polymerase gene. The Maximum-likelihood tree was calculated by using the MEGA X software. Statistical support for nodes was obtained by bootstrapping (1,000 replicates); only values ≥50% are shown. Virus names are preceded by the respective NCBI GenBank accession number. Sequences generated during this study are marked in red. The chart background of viruses belonging to the same coronavirus genus is highlighted by the same color and the genera are indicated.

To further confirm that there is no ongoing virus circulation in the sampled rat population, we tested the lungs by a SARS-CoV-2-specific real-time RT-PCR targeting the RNA-dependent RNA polymerase (RdRp) gene *(9)* and by a likewise RdRp-based, generic pan-coronavirus RT-PCR *(10)*. In the SARS-CoV-2-specific test, all samples scored negative, verifying the absence of SARS-CoV-2 in the analyzed samples from Berlin. Nevertheless, five lung samples were positive in the pan-coronavirus RT-PCR; all five animals were trapped at the same location (Figure 1). For further characterization, the RT-PCR products were sequenced in both directions with the primers used for amplification. The amplicon sequences (NCBI GenBank accession numbers OR854629-OR854633) were subsequently compared to representative coronavirus sequences obtained from GenBank. Virus typing based on the partial RdRp sequences revealed that the viruses found in Berlin rats belong to the genus *Alphacoronavirus* and are closely related to each other (99.4-100.0% identity on nucleotide level) and to the Lucheng Rn rat coronavirus (Figure 1B). Hence, in contrast to SARS-CoV-2, rodent-associated alphacoronaviruses appear to circulate in the investigated rat population, which is in line with previous studies investigating coronaviruses in rats *(2,5)*.

Viral monitoring of rodent populations like rats is essential to understand e.g. virus occurrence, transmission characteristics and pathogenesis, not only for their potential impact on rodents but also due to the potential for recombination and the zoonotic nature of coronaviruses. Research into rodent coronaviruses contributes to a broader understanding of these viruses and aids in the development of strategies for managing both animal and public health.

## Acknowledgments

We thank Bianka Hillmann, Dennis Karnatz and Janina Beyer for excellent technical assistance, our colleagues from the FLI for the help during dissection and the pest controllers from Berlin for providing rats from rodent pest control. The study was supported by intramural funding of the German Federal Ministry of Food and Agriculture provided to the Friedrich-Loeffler-Institut, partial funding from the European Union Horizon 2020 project (Versatile Emerging infectious disease Observatory, grant no. 874735; awarded to M.B.) and DZIF, thematic translational unit (TTU) “Emerging Infections” (awarded to R.G.U.).

## Ethical Statement

The rat samples were collected in the context of rodent pest control, which did not require a specific permit.

## Notes

### Competing Interest Statement

The authors have declared no competing interest.

